# Are indigenous territories effective natural climate solutions? A neotropical analysis using matching methods and geographic discontinuity designs

**DOI:** 10.1101/2020.12.23.424126

**Authors:** Camilo Alejo, Chris Meyer, Wayne S. Walker, Seth R. Gorelik, Carmen Josse, Jose Luis Aragon-Osejo, Sandra Rios, Cicero Augusto, Andres Llanos, Oliver T. Coomes, Catherine Potvin

## Abstract

Indigenous Territories (ITs) with less centralized forest governance than Protected Areas (PAs) may represent cost-effective natural climate solutions to meet the Paris agreement. However, the literature has been limited to examining the effect of ITs on deforestation, despite the influence of anthropogenic degradation. Thus, little is known about the temporal and spatial effect of allocating ITs on carbon stocks dynamics that account for losses from deforestation and degradation. Using Amazon Basin countries and Panama, this study aims to estimate the temporal and spatial effects of ITs and PAs on carbon stocks. To estimate the temporal effects, we use annual carbon density maps, matching analysis, and linear mixed models. Furthermore, we explore the spatial heterogeneity of these estimates through geographic discontinuity designs, allowing us to assess the spatial effect of ITs and PAs boundaries on carbon stocks. The temporal effects highlight that allocating ITs preserves carbon stocks and buffer losses as PAs in Panama and Amazon Basin countries. The geographic discontinuity designs reveal that ITs’ boundaries secure more extensive carbon stocks than their surroundings, and this difference tends to increase towards the least accessible areas, suggesting that indigenous land use in neotropical forests may have a temporarily and spatially stable impact on carbon stocks. Our findings imply that ITs in neotropical forests support Nationally Determined Contributions (NDCs) under the Paris Agreement. Thus, Indigenous peoples must become recipients of countries’ results-based payments.

## Introduction

Avoided forest conversion and natural forest management are among the most cost-effective natural climate solutions to meet the Paris Agreement (1). Protected Areas (PAs), cornerstones of biodiversity conservation, may contribute to these cost-effective solutions by preventing carbon stocks losses (2). However, since 1990, South America and Central America have tripled the area of PAs (3) while simultaneously losing 10% and 25% of forest cover, respectively (4). These forest conversion trends stress the need for additional natural climate solutions that could reinforce the role of PAs. In Neotropical countries and across the globe, Indigenous Territories (ITs) cover significant portions of natural lands with minimal human disturbance and tend to overlap with PAs (5). More than 30% of the Amazon Basin forest’s aboveground carbon stocks are in ITs, and nearly 7% of these stocks are in areas overlapping with PAs (Overlapped Areas, hereafter OAs) (6). Thus, ITs and OAs with less centralized governance and providing livelihoods may conserve forests and potentially represent effective natural climate solutions.

However, the effect of ITs, OAs and PAs in forest conservation might be overestimated. These land tenures tend to be located in higher elevations, steeper slopes, and greater distances to roads and cities than unprotected lands, lowering deforestation probabilities (7,8). To control for this non-random spatial location, an increasing number of studies have relied on a statistical technique called matching analysis (9,10). In these studies, matching analysis samples observations with similar geographical characteristics, removing heterogeneous observations, and allowing to compare ITs, OAs, and PAs with unprotected lands. For example, using matching analysis, ITs in the Brazilian Amazon have been found to restrain high deforestation pressure more effectively than PAs (11). Panama’s PAs and untitled ITs more effectively avoided deforestation than unprotected lands with similar topography and accessibility (12). Matching analysis also allowed identifying decreased deforestation where ITs and other land tenures overlap (e.g., PAs) in Peru (Anderson et al., 2018). Furthermore, Blackman & Veit (2018) concluded that ITs in the Amazon Basin of Colombia, Bolivia and Brazil avoid carbon emissions from deforestation. Therefore, controlling for spatial location using matching supports the claim that ITs are as effective as PAs to avoid deforestation.

Despite the influence of anthropogenic degradation and recovery on forest conservation and carbon stocks dynamics, research on matching analysis has been limited to examining the effect of land tenures on avoided deforestation. Shifting cultivation, considered a driver of degradation (15), is common among tropical forest landholders (16). After long fallow periods (>20 years), shifting cultivation can only recover around 50% of mature forests’ carbon stocks (17). Logging and fires, other causes of degradation in tropical forests, remove 45% and 22% of forest’s carbon stocks and take decades to recover (18). Thus, accounting for forest degradation and recovery in temporal carbon stocks dynamics may shed a different light on the effectiveness of land tenures in forest conservation, particularly in those with fewer use restrictions (e.g., ITs and OAs). However, little is known about the temporal effect of ITs, OAs, and PAs on carbon stocks dynamics after controlling for spatial location.

Matching analysis controls for spatial location, but it does not guarantee unambiguous estimates of land tenure effects in forest conservation. Karsenty et al. (19) highlight that matching analysis implies weighting influence to particular deforestation (or degradation) covariates, such as roads or rivers. The choice and omission of covariates influence the observations sampled by matching, potentially excluding relevant areas, and altering the effect attributed to a particular land tenure (19). In this regard, some have recognized that sampling through matching analysis might not be independent and exclude observations around the boundaries of protected lands (20–22), rather than exploring the implications of sampling across these boundaries. Conversely, the effect of ITs and PAs boundaries on deforestation has been estimated through regression discontinuity designs. Bonilla-Mejía & Higuera-Mendieta (23) found that ITs’ boundaries are more effective than PAs at curbing deforestation in Colombia. Similarly, Baragwanath & Bayi (24) established that ITs’ boundaries with granted property rights in Brazil decrease deforestation. However, few studies have used matching analysis in geographic discontinuity designs, control for geographic distance among observations (25), and estimate the effect of ITs and PAs boundaries on carbon stocks. Nor have they addressed whether land tenures with different forest governance, such as ITs and PAs, imply different spatial effects on carbon stocks.

This study builds upon previous research assessing the effect of land tenures on deforestation through matching analysis and addresses some limitations of this methodology. Using Panama and Amazon Basin Countries, this study aims to estimate ITs, OAs, and PAs temporal and spatial effects on aboveground carbon stocks. The hypothesis is that PAs with centralized governance and disincentives on forest use will secure higher carbon stocks than ITs and OAs over time and throughout their boundaries by reducing the influence of anthropogenic degradation. Regardless of forest use disincentives and governance, we find that PAs, OAs, and ITs preserve carbon stocks and buffer losses temporarily and spatially across neotropical forests.

Our study makes three contributions to the literature. First, we provide a consistent use of matching analysis in multiple land tenures and countries, allowing us to compare the effects of ITs, OAs, and PAs across neotropical forests. Conversely, previous studies have analyzed either multiple land tenures on a country scale (e.g., 14,17,18) or single land tenure categories across regions (e.g., 11,16). Second, we use the temporal dynamics of aboveground carbon stocks (2003 to 2016) instead of forest cover, thus making it possible to estimate a more accurate temporal effect of ITs, OAs, and PAs in climate change mitigation. Furthermore, we explore the spatial heterogeneity of these effects through geographic discontinuity designs, allowing us to assess the spatial effect of ITs, OAs, and PAs boundaries on carbon stocks. To our knowledge, this study is among the first to estimate the effect of multiple land tenures on carbon stocks temporarily (14 years) and spatially (throughout boundaries), providing a quantified estimate of forest conservation and climate change mitigation across Neotropical Forests.

### Theory of change

Our study assumes a causal relationship between ITs, OAs and PAs, the treatments, and forest’s carbon stocks, the outcome. Here, we explain the different components and assumptions for this causal relation to occur (Fig 1). Spatial location covariates influencing the suitability of agriculture (e.g., altitude and slope) and market pressure (human settlements, roads, rivers) (28,29) represent input components driving carbon stocks losses in the treatments and controls (other lands). ITs, OAs, and PAs are known to experience an overall reduced influence from these covariates compared with other lands (7). Moreover, as market pressure declines inside the treatments boundaries (30), forest cover increases (31). Beyond the influence from these covariates, we expect ITs, OAs, and PAs to directly cause positive outcomes, that is, securing larger carbon stocks than other lands (the control). However, these land tenures, are subject to external and indigenous governance (14,32) that may result in different outcomes.

**Fig 1.**
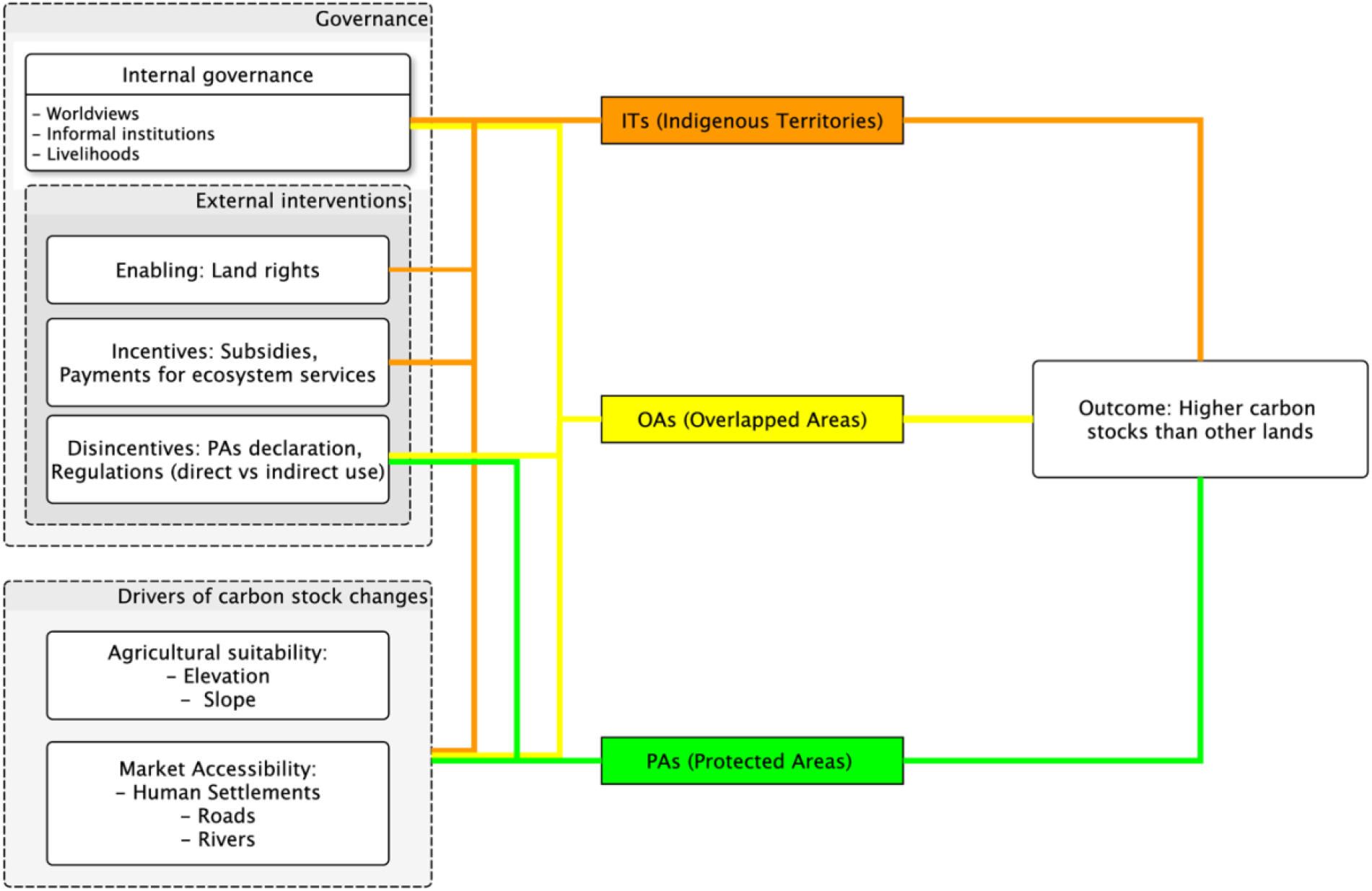
A generic theory of change for land tenures effects on carbon stocks. The lines symbolize hypothetical pathways of how governance components and drivers of carbon stocks change influence ITs (orange), OAs (yellow), PAs (green) to result in an expected outcome.

ITs may result in positive outcomes due to indigenous and external governance. Indigenous governance emerges from worldviews and cultural values that do not privilege ecosystem conservation at the expense of local livelihoods or vice-versa (33,34). These forms of governance build informal institutions that restrict access to other agents and limit the spatial and temporal extent of agriculture and other livelihood activities (35). Thus, even if deforestation and degradation caused by permanent and shifting agriculture, logging, and firewood extraction reduce forests’ carbon stocks (36), their negative effect is expected to be temporarily and spatially limited in ITs compared with other lands. Furthermore, external governance interventions may limit the influence of local livelihoods on forests (37). For example, governments’ recognition of land rights (38) or incentives that reward communities for forest conservation actions may contribute to secure carbon stocks (39,40).

Regarding PAs and OAs, we assume a predominant influence of external governance. The declaration of PAs (in public or private lands) represents government disincentives to restrict land use, conserve forests (32), and consequently limit carbon stock losses temporarily and spatially. While certain government regulations may allow direct uses to some agents, PAs tend to have centralized forest governance (41). OAs are PAs established in ITs and have been interpreted as external interventions that privilege conservation and limit indigenous governance and livelihoods (42). Consequently, OAs represent an intermediate treatment between ITs and PAs that also result in limited carbon stock losses compared with other lands. Given that PAs constitute the highest limitation on forest livelihoods, and therefore deforestation, and degradation, we expect that they will result in more substantial effects on carbon stocks than OAs and ITs.

## Methods

### Geographic scope

The ideas developed in this study emerged from discussions during the annual meeting of the “Red Amazónica de Información Socioambiental Georeferenciada” RAISG (Amazon Georeferenced Socio-Environmental Information Network) carried out in Quito (Ecuador) in August 2018. The authors belong to diverse organizations that participate or collaborate with RAISG. Additionally, some of the authors have collaborated with the “Coordinadora de las Organizaciones Indígenas de la Cuenca Amazónica” - COICA (Coordinator of Indigenous Organizations of the Amazon River Basin), which also participates in RAISG, and the “Alianza Mesoamericana de Pueblos y Bosques” - AMPB (Mesoamerican Alliance of Peoples and Forests). Regarding this study, these collaborations have resulted in sharing and curating geospatial information on PAs and ITs that define our study’s geographical scope: Panama and the Amazon Basin portions from Colombia, Ecuador, Peru, and Brazil. Only the authors participated in the research design and the interpretation of the results.

Our study focuses on three land tenures in Panama and Amazon Basin Countries (Fig 2): PAs, ITs, and OAs. PAs encompass national and subnational jurisdictions with governance by governments, private governance, and shared governance that allow sustainable use from privates and communities (Table 1). ITs without official titles or in the process of official recognition (i.e., untitled lands) were also included, except in Colombia, where the data was not available. All ITs overlapping with PAs were defined as OAs. All private and public lands outside ITs, OAs and PAs were defined as other lands.

**Fig 2.**
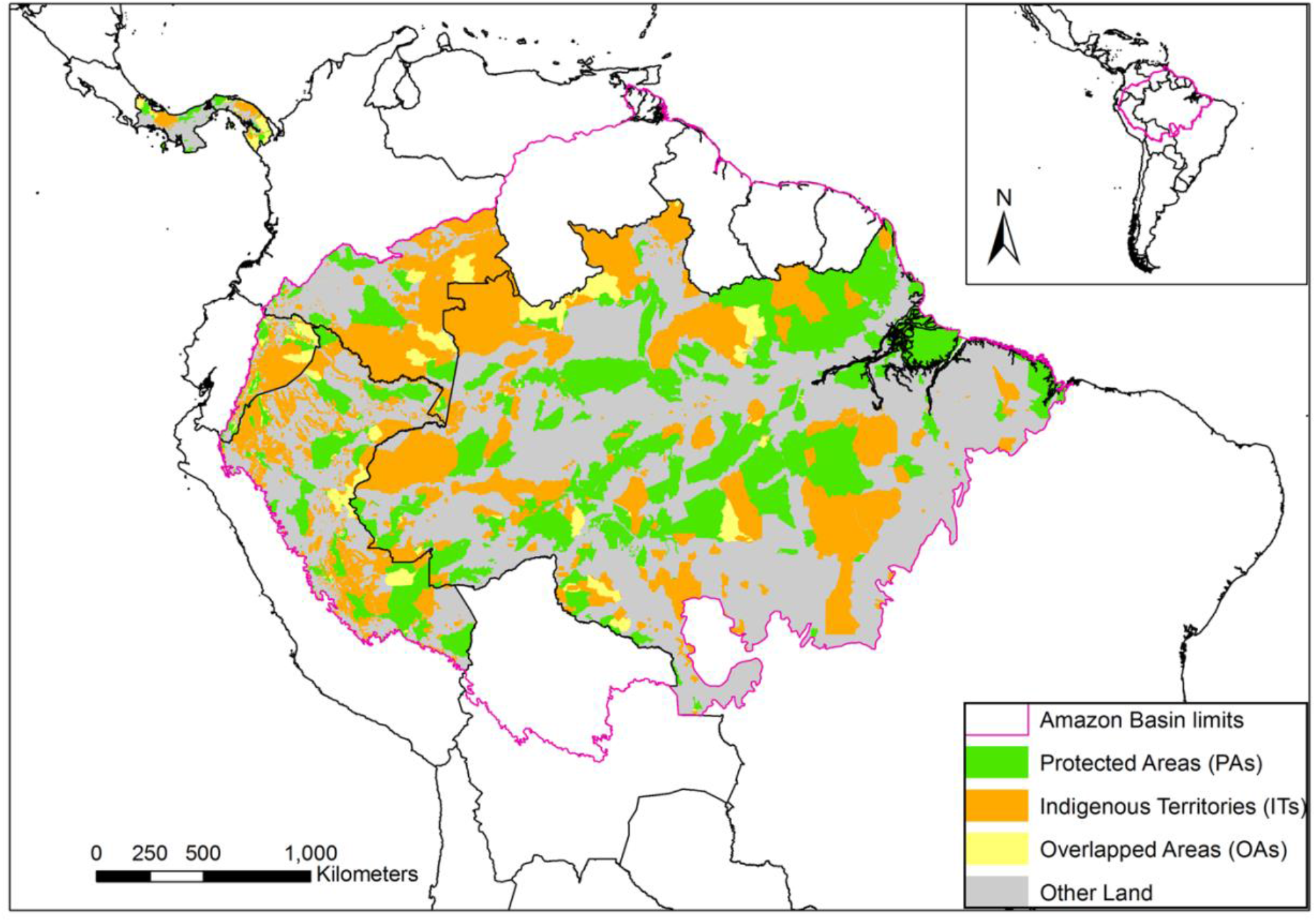
Study Area. Panama and the Amazon Basin portions of Colombia, Ecuador, Peru, and Brazil. Land tenure is classified as PAs (green), ITs (orange), OAs (yellow), and Other Land (grey).

**Table 1.** PAs and ITs included in the study.

| Country | Protected Areas (PAs) | PAs IUCN Category | Indigenous Territories (ITs) |
| --- | --- | --- | --- |
| Panama | National Park | II | Titled: "Comarcas" |
|  | Protective Forest | V | Titled: Collective Territories |
|  | Wildlife Refugee | IV | Claimed/Untitled |
|  | Multiple Use Area | VI |  |
|  | Forest Reserve | IV |  |
|  | Hydrological Reserve | V |  |
|  | Zone of hydrological protection | V |  |
| Colombia | National Park | I-II | Titled: Indigenous Reserve |
|  | National Protective Forest Reserve | VI |  |
|  | National Forest Reserve | I |  |
|  | Civil Society Nature Reserve | VI |  |
|  | Fauna and Flora Sanctuary | IV |  |
| Ecuador | National Park | NR* | Titled |
|  | Protective Forests | NR | Declared |
|  | Ecological Conservation Area | NR* | Claimed/Untitlled |
|  | Biological Reserve | NR* |  |
|  | Ecological Reserve | NR* |  |
|  | Fauna Production Reserve | NR |  |
|  | Wildlife Refugees | NR |  |
| Peru | National Park | NR* | Titled / Declared: Native community |
|  | National Sanctuary | NR* | Titled/ Declared: Peasant community |
|  | Historical Sanctuary | NR* | Claimed/Untitlled |
|  | Protective Forest | NR |  |
|  | Landscape Reserve | NR |  |
|  | Communal Reserve | NR |  |
|  | Hunting Reserve | NR |  |
| Brazil | National Park | II - NR* | Titled/ Declared: Indigenous Area |
|  | Environmental Protection Area | NR | Titled/ Declared: Native Community |
|  | Area of Relevant Ecological Interest | NR | Titled / Declared: Indigenous Reserve |
|  | Ecological Station | NR* | Titled / Declared: Indigenous Territory |
|  | Natural Monument | III | Claimed/Untitlled |
|  | Nature Reserve | NR* |  |
|  | Biological Reserve | NR* |  |
|  | Sustainable Use Reserve | NR |  |
|  | Ecological Reserve | NR* |  |
|  | Extractive Reserve | NR |  |
|  | State Forest | NR* |  |
|  | State Park | NR* |  |
|  | Wildlife Refugee | NR* |  |

PAs are accompanied by IUCN categories, except when not reported (NR) and only allowing indirect use*. ITs overlapping with PAs are considered OAs.

### Spatial data and processing

The boundaries of ITs and PAs were curated by the Neotropical Ecology Laboratory (McGill University, Smithsonian Tropical Research Institute) for Panama; and RAISG (Amazon Geo-referenced Socio-Environmental Information Network) in the case of Amazon Basin Countries. This spatial information was used to determine the overlaps of ITs and PAs, here defined as OAs.

We used Annual carbon density maps based on raster data (∼500 m resolution) that was generated by the Woodwell Climate Research Centre between 2003 and 2016 and explained in detail by Baccini et al. (43,44) and Walker et al. (6). These estimations derive from combining LiDAR data and field measurements that calibrate a machine learning algorithm that generates annual carbon density estimates from MODIS satellite imagery. These carbon density maps can detect annual losses and gains in carbon density, aggregating changes from deforestation, forest degradation, and recovery.

Elevation, slope and the distance to roads, settlements and rivers were included as covariates to establish the spatial location conditions associated with annual carbon density across countries (S1A Appendix). Elevation and slope were obtained from the satellite imagery of the SRTM (Shuttle Radar Topographic Mission - Arc Second Global). The distance to roads was calculated from geospatial data produced by national institutions in Panama. Road distance corresponding to Amazon Basin countries was based on the geospatial data curated by RAISG. The distances to rivers and settlements (> 5000 people) were calculated from geospatial data produced by national institutions. Land tenure and covariate data were resampled to the spatial resolution of carbon density, creating observation units of ∼500-m resolution across different land tenures with estimates for covariates and carbon density. All geoprocessing was performed in ArcGIS (ESRI, 2018). Finally, we established the non-random spatial location of ITs, OAs, and PAs by estimating their mean covariate differences with other lands in each study area using Mann Whitney tests (S1B Appendix).

### Temporal effects on carbon stocks

As an initial analysis, we performed matching analysis and linear mixed models to control for spatial location and infer the temporal effect of ITs, OAs, and PAs on carbon stocks relative to other lands (Fig 3). Matching analysis preprocesses datasets to reduce the association of a treatment variable with covariates by removing heterogeneous observations and creating a subset of treatment and control observation units with similar covariate values (45). Here, the treatment variable corresponded to land tenure, and matching created subsets of observation units of ∼500 m resolution in the treatment (i.e., ITs, OAs, and PAs) and control (i.e., other lands) with similar slope, elevation, and distance to roads, settlements, and rivers. To account for the size and heterogeneity of the Brazilian Amazon, we included the states as covariates in this country.

**Fig 3.**
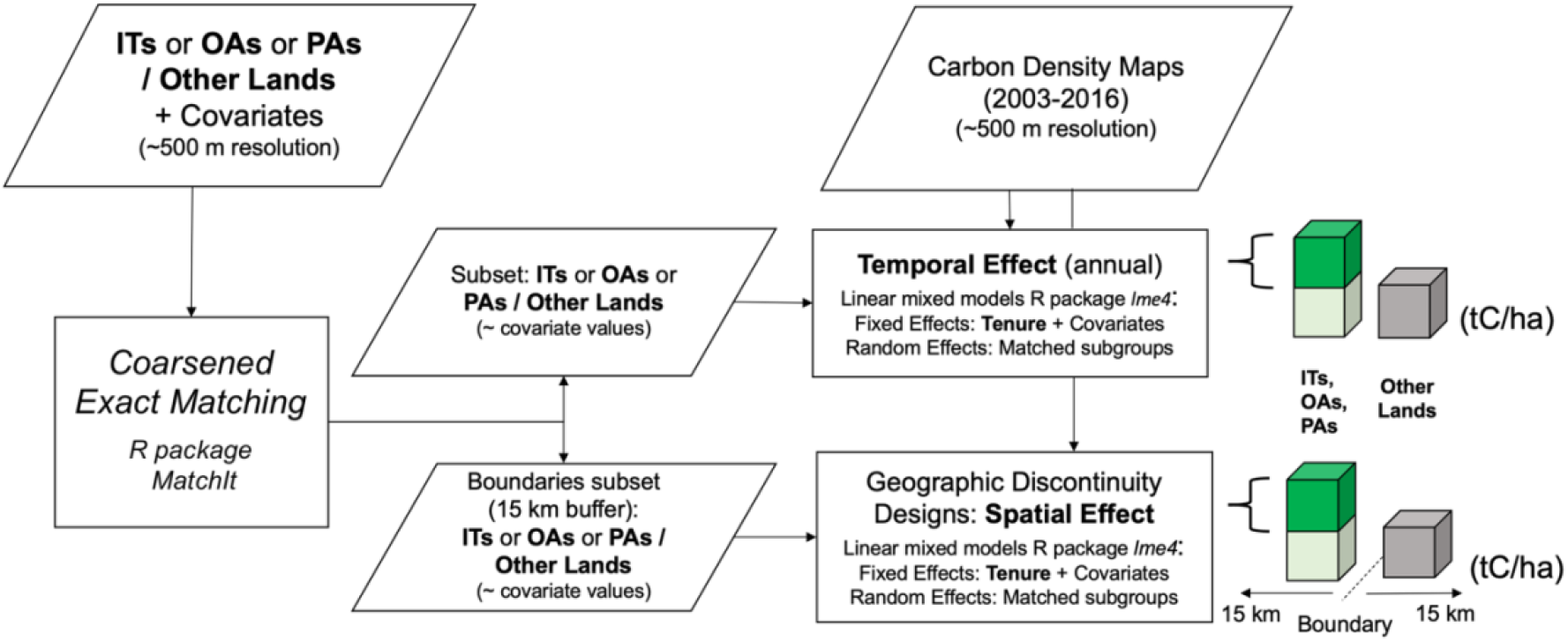
Workflow to infer the temporal and spatial effect of ITs, OAs, and PAs on carbon stocks.

Specifically, we used coarsened exact matching (CEM) (46) with the R package *MatchIt* (47) for ITs, OAs, and PAs in all study areas. Following steps from Iacus et al. (48), we first defined coarsening choices for each covariate (S1C Appendix). For example, the elevation was coarsened in multiple categories based on 100 meters intervals. This coarsening choice meant that observation units with elevation values between 900 and 1000 m were considered “equivalent”. Then, CEM located control and treatment observation units in matching sub-groups with equivalent coarsened values for all covariates. The third step pruned matching sub-groups that did not have at least one treatment and one control observation with equivalent coarsened covariate values. These steps were reiterative until the coarsening choices produced a covariate balance between treatments and controls. The covariate balance before and after matching was assessed through standardized mean differences and Kolmogorov-Smirnov statistics (49) (S2A and S2B Appendices). The balance assessments were performed in the R package *Cobalt* (50).

After isolating the effect of spatial location through matching, we made temporal estimates regarding the effect of allocating ITs, OAs, and PAs on carbon stocks in each country. This effect was calculated using linear mixed models in the R package *lme4* (51) with the general expression defined as:

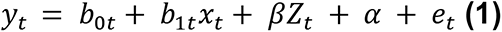

where *y_t_* was carbon density in year *t*, the outcome variable, and *b_0t_* was the fixed intercept. b_1t_ and *x_t_* were the fixed effect slope and predictor of land tenure (i.e., dummy for ITs, OAs, and PAs), respectively. Additionally, β was a vector of additional fixed effects for a vector of predictors Z_t_, containing the covariates elevation, slope, and distance to roads, settlements, and rivers. Including the covariates as fixed effects span any remaining imbalances from the matched subsets. The matched sub-group (matched observation units in treatments and control with similar covariate values) was the random effect α_t_ to account for the structure of the matched subsets. These linear mixed models were estimated annually between 2003 and 2016 in all ITs, OAs, and PAs and study areas. Two parameters derived from the linear mixed models were used to determine the effect of ITs, OAs, and PAs on carbon stocks after controlling for spatial location: the fixed effects intercept *b_ot_* and fixed effects slope *b_1t_*. *b_ot_* refers to the average annual carbon density found in other lands lacking a protected status and represents the carbon stocks baseline for ITs, OAs, and PAs. *b_1t_* refers to the annual average differences of carbon stocks between these land tenures and other lands, defined as the temporal effect.

### Spatial effects on carbon stocks

After calculating the distance of matched observation units around the boundaries of ITs, OAs, and PAs, we used geographic discontinuity designs to examine the spatial heterogeneity of the temporal effects. Geographic discontinuity designs estimate the effect of administrative boundaries (52), here defined as spatial effects. Specifically, we assessed how ITs, OAs, and PAs boundaries influence carbon stocks compared with other neighbouring lands. Our geographic discontinuity designs are based on two assumptions. First, following Keele et al. (25), we assume that after controlling for covariates and the geographic distance (i.e., the distance among observations throughout a boundary), the treatment assignment occurs as-if randomized, allowing to estimate the spatial effects. Our second assumption, which derives from the first, is that the spatial effect is a function of the treatment of interest (52). This assumption implies that the boundaries of ITs, OAs, and PAs will influence carbon stocks.

To implement the geographic discontinuity designs, we created subsets of observation units with buffer zones inside and outside of ITs, OAs, and PAs boundaries of 0-0.5 km, 0–1 km, 0–5 km, 0-10 km, and 0–15 km. We chose these buffer zones because covariates’ pressure usually ceases between 5 and 10 km (30), and the vegetation seems to stabilize in PAs and ITs around 15 km (31). Similar to Keele et al. (25), we used matching methods to find treatment and control observation units with similar covariates. As the temporal effects matching, we performed CEM within the buffer zones subsets, including slope, elevation, and distance to roads, settlements, and rivers as covariates. Additionally, we controlled for the geographic distance among observation units according to buffer zones. For example, in buffer zones 0–1 km, we included matches across a 2-km radius, and in 0–15-km buffer zones, a 30-km radius. The covariate balance before and after matching was assessed through standardized mean differences and Kolmogorov-Smirnov statistics (S2C and S2D Appendices).

The differences between average carbon stocks stored inside and outside the boundaries of ITs, OAs, and PAs, or the spatial effects, were also estimated through the linear mixed models aforementioned in 2003 and 2016. The covariates were included as fixed effects, spanning any remaining imbalances from matching. To support the credibility of the spatial effects, we performed falsification tests where each covariate in Z_t_ was treated as an outcome variable *y_t_* according to the linear mixed model above (52). The falsification tests showed that allocating ITs, OAs, and PAs had negligible effects on the covariates after matching (S2E Appendix). The annual spatial effects, covariate balance tests, and falsification tests were estimated in all ITs, OAs, and PAs, across multiple buffer zones (0-1 to 0-15 km) and study areas.

The geographic discontinuity designs support the previous assumptions. Matching guarantees that observations inside and outside ITs, OAs, and PAs will be valid counterfactuals as they share distance to the boundaries (e.g., 0-1 km), mutual proximity (e.g., 2 km radius), and covariates influence. This role of matching is confirmed by the covariate balance tests and the falsification tests. Thus, the treatments assignment (i.e., ITs, OAs, and PAs) occur as-if randomized (first assumption). Moreover, we account for local effects by matching valid counterfactuals in neighboring subgroups and incorporating them as random effects in the linear mixed models. These local effects might control the influence of unobserved covariates that operate on a local or restricted geographical scale, ensuring that the overall spatial effect is a function of the treatments (second assumption). Finally, if the assumption of valid counterfactuals holds across multiple buffer distances to treatments boundaries, it is possible to estimate the heterogeneity of the spatial effects. In other words, it is possible to explore how the spatial effects vary at multiple distances from ITs’, OAs’, and PAs’ boundaries.

### Sensitivity analysis

Matching is expected to control for observed covariates and correlated unmeasured covariates (49). In our study, unmeasured covariates that influence carbon stocks include population, opportunity costs from agriculture and cattle, or the probability of fire occurrence. However, these unmeasured covariates are correlated with other covariates of market pressure and agricultural suitability (9,53) (i.e., the observed covariates). Thus, we used sensitivity analyses to assess the effect of unmeasured covariates unrelated to the observed covariates but related to the treatments (i.e., ITs, OAs, and PAs) and their effects (temporal or spatial) (54).

Particularly, we estimated the E-value, which represents the minimum strength that an unmeasured covariate would need to have with the treatment and its effect, for the treatment and effect association not to be causal (55). This value is an estimate that accommodates effects from observational studies (i.e., not randomly assigning the treatment and control) that do not have 1-1 matched pairs (CEM matches multiple observations in subgroups) (54). The E-value can be calculated by the expressions:

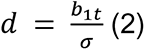

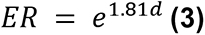

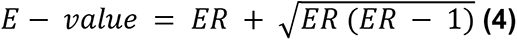

where *b_1t_* is the fixed effects slope for land tenure, that is the temporal or spatial effect, σ is the standard deviation of the temporal or spatial effect, *d* is the standardized temporal or spatial effect, and ER is the effect ratio. ER, equivalent to a Risk Ratio, compares the probability of a positive spatial/temporal effect in ITs, OAs, and PAs with the probability of a positive effect in other lands. The expressions above are further justified in (55). In our study, an ER greater than 1 indicates a greater probability that ITs or OAs, or PAs will store higher carbon stocks than other lands, either temporarily or spatially. For example, an ER of 2 in ITs means that, after controlling for covariates, ITs are two times more likely to store higher carbon stocks than other lands. A hypothetical E-value of 3 would imply that the ER of 2 could be explained away by an unmeasured covariate that was associated with both the allocation of ITs (i.e., treatment) and annual carbon stocks (i.e., outcome) each by 3-fold, above and beyond the observed covariates. However, a weaker unmeasured confounding could not alter the ER and, therefore, the spatial and temporal effects. Following (56), the E-value assesses the strength of an unmeasured covariate to alter the temporal and spatial effect of ITs, OAs, and PAs on carbon stocks. This value was estimated with the R package *Evalue* (57) for all spatial and temporal effects in 2003 and 2016 across the study areas.

## Results

### The temporal effect of Indigenous Territories and Protected Areas on carbon stocks

Matching analysis and the linear mixed models controlled the influence of spatial location covariates, allowing to estimate the temporal effect of allocating ITs, OAs, and PAs on carbon stocks. This temporal effect represents the annual mean difference of carbon stocks between these land tenures and other lands. Across Panama and Amazon Basin countries, the carbon stocks from 2003 to 2016 in ITs, OAs, and PAs were usually higher than other lands (i.e., the baseline), resulting in positive temporal effects (Fig 4). According to sensitivity analyses, an unmeasured covariate would need to have a stronger effect than ITs, OAs, and PAs through pathways independent of the covariates to modify these temporal effects (S3A Appendix).

**Fig 4.**
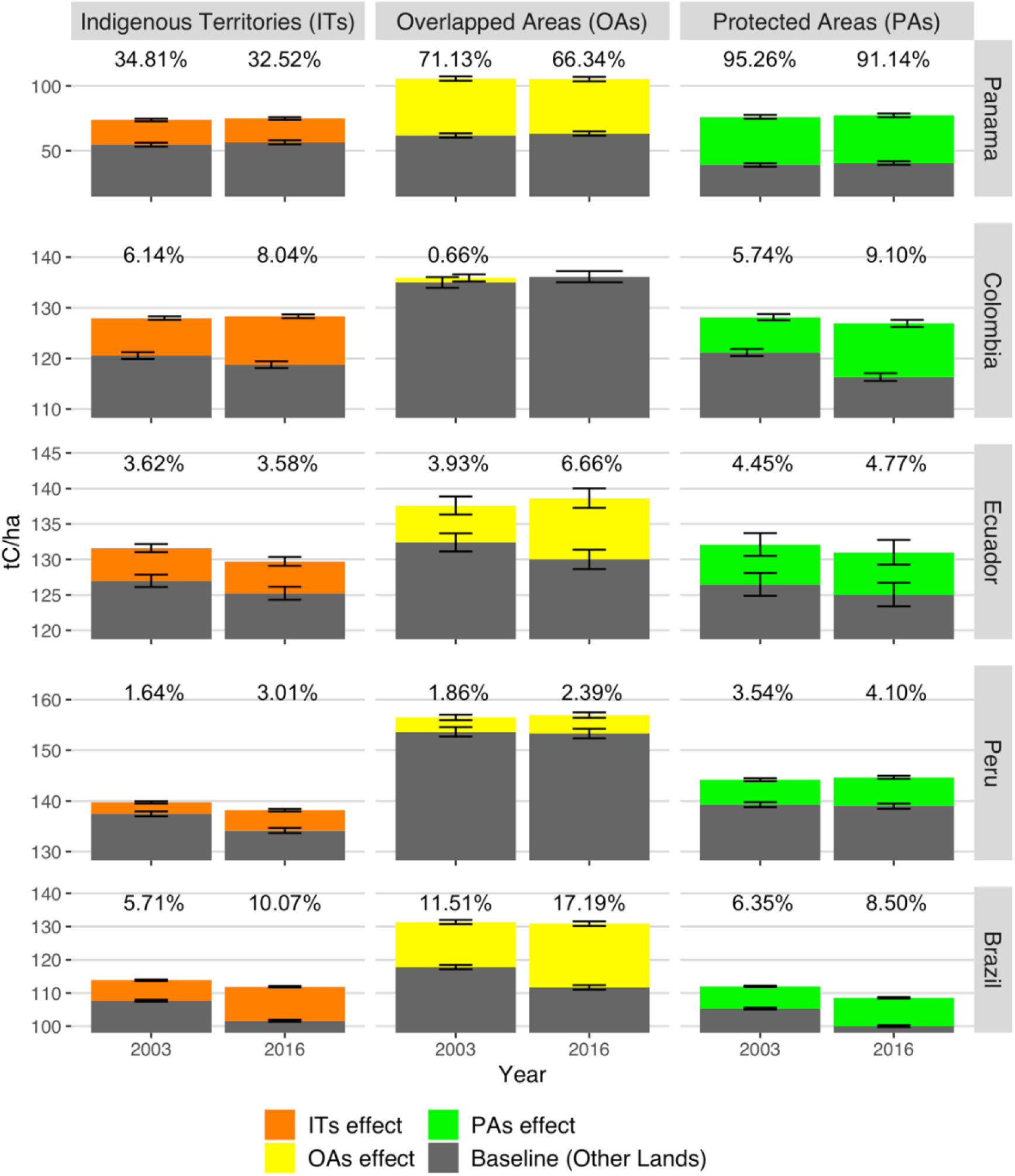
The temporal effects of ITs, OAs and PAs on aboveground carbon stocks across neotropical countries in 2003 and 2016. Significant temporal effects (p < 0.05) are represented as colored bars and percentages, indicating the additional/fewer carbon stocks secured by allocating ITs (orange), OAs (yellow), and PAs (green) relative to the baseline (Other Lands, grey) after controlling for spatial location. Error bars reflect 95% confidence intervals for the baselines and temporal effects.

Country-level comparisons of temporal effects in ITs, OAs, and PAs reveal three regional patterns (Fig 4). Panama had low carbon stocks baselines in other lands (< 65 t C/ha) and substantial temporal effects that represented an increase in carbon stocks above 30%. Brazil displayed moderate baselines (< 115 t C/ha) and temporal effects (< 18%). The carbon stocks baselines in western Amazon Basin countries exceeded those of Brazil (> 115 t C/ha), while the temporal effects were moderate (< 10%). Hence, the temporal effects seem substantial in countries with reduced carbon stocks in other lands.

The positive temporal effects also reveal the additional amount of carbon stocks secured by allocating ITs, OAs, and PAs in a particular year compared to other lands (i.e., baseline) across Panama and Amazon Basin countries (Fig 4). During 2003, PAs in Panama secured 95% (37 t C/ha) larger carbon stocks than their baseline (39 t C/ha). Relative to more substantial baselines (> 55 t C/ha), Panama’s ITs and OAs accounted for 35% (19 t C/ha) and 71% (44 t C/ha) additional carbon stocks. Similar to Panama, ITs, OAs, and PAs in Amazon Basin countries represented positive temporal effects in 2003. Brazil’s ITs and PAs represented 6% (∼6 t C/ha, respectively) additional carbon stocks compared to their baselines (∼105 t C/ha), and this effect nearly doubled in OAs (12%, 14 t C/ha). Western Amazon Basin countries displayed similar temporal effects in 2003, ranging between 1.6 – 6.1% (i.e., 5 - 7 t C/ha) in ITs from Peru and Colombia, 3.5 – 5.7 % (i.e., 5 - 7 t C/ha) in PAs from the same countries, and 0.7 – 4 % (i.e., 0.5 - 5 t C/ha) in OAs from Colombia and Ecuador. Despite regional differences, these results suggest that in 2003 OAs and ITs had a similar effect on carbon stocks compared to PAs in neotropical countries.

Overall, the temporal effects on carbon stocks remained stable or increased relative to other lands until 2016 (Fig 4, S4A Appendix). These effects remained stable in PAs and ITs from Ecuador and did not vary more than 0.5%. ITs in other Amazon Basin countries exhibited increases in temporal effects, reaching between ∼ 3% (4 t C/ha) in Peru and ∼10% (10 t C/ha) in Brazil. Similarly, Amazon basin PAs had increases that resulted in temporal effects between ∼ 4% (∼11 t C/ha) and ∼ 9.1% (9.5 t C/ha) for Peru and Colombia, respectively. The temporal effects considerably varied in Amazon Basin OAs during 2016, showing no differences with the baseline in Colombia and the largest increase in Brazil (17.2%, 19 t C/ha). Conversely, ITs, OAs, and PAs in Panama experienced decreases in temporal effects (> −5%) that seem to be driven by the recovery of carbon stocks in other lands (S4B Appendix). Thus, stable and increasing temporal effects reflect that allocating ITs, OAs, and PAs buffered losses and secured the stability of carbon stocks relative to the other lands. Furthermore, these results reveal that indigenous lands (i.e., ITs and OAs) and PAs secured similar amounts of carbon stocks until 2016.

### Insight at a finer scale: the spatial effect of Indigenous Territories and Protected Areas on carbon stocks

To identify the spatial implications of matching analysis in quantifying forest conservation, we estimated the distance of observation units to the boundaries of ITs, OAs, and PAs (Fig 5, Table 2). Matched observation units in these land tenures had a range of average distances to their boundaries, between 1.3 km (± 2.26) in PAs from Ecuador and 10.15 km (± 11.70) in PAs from Peru. The distance of matched observation units in other lands to ITs’, OAs’, and PAs’ boundaries ranged between 3.10 km (± 3.13) (Ecuador) and 9.52 km (± 7.72) (Panama). Not surprisingly, the spatial distributions imply that observations along these boundaries are more likely to share spatial features (i.e., elevation, slope, and distance to roads, settlements, and rivers). Moreover, most human settlements inside ITs, OAs, and PAs tend to be located in these accessible areas, especially less than 5 km from the boundaries (S4C Appendix). In the case of observations in ITs, OAs, and PAs, these sampling outcomes suggest that matching analysis selects the most accessible areas, omitting the core and possibly more intact forests. Thus, the spatial distribution from matching indicates that ITs’, OAs’, and PAs’ temporal effects are conservative.

**Fig 5.**
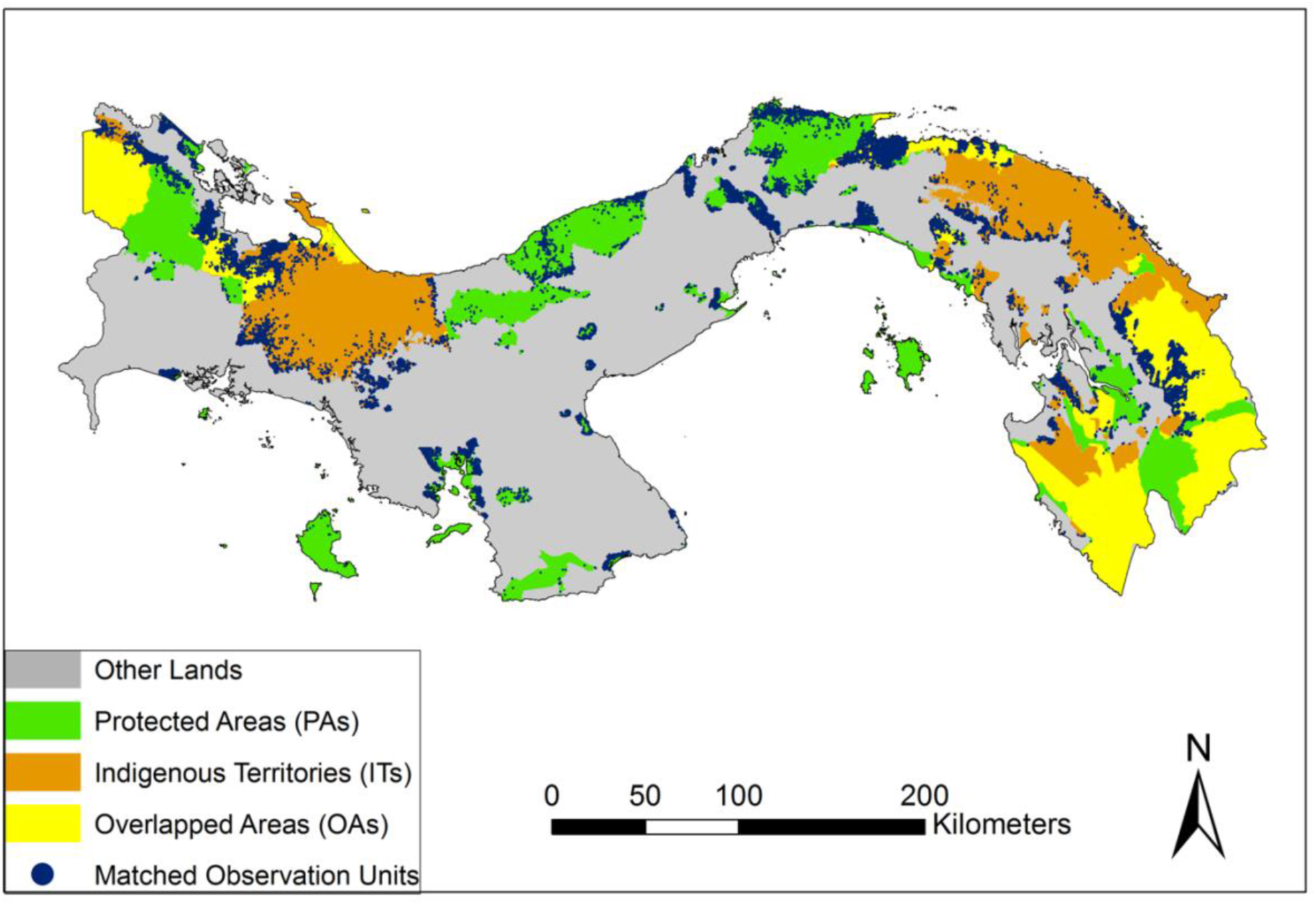
Observation units sampled through matching analysis in ITs, OAs, and PAs from Panama.

**Table 2.** Mean distance to ITs’, OAs’, and PAs’ boundaries of observation units sampled through matching analysis by country and land tenure.

| Country | Land tenure | Mean distance to boundaries (km) | SD |
| --- | --- | --- | --- |
| Panama | Other Lands | 9.51 | 7.72 |
|  | PAs | 1.04 | 1.41 |
|  | ITs | 2.37 | 2.99 |
|  | OAs | 2.25 | 2.75 |
| Colombia | Other Lands | 10.57 | 10.70 |
|  | PAs | 6.32 | 5.25 |
|  | ITs | 9.35 | 1.34 |
|  | OAs | 8.69 | 7.55 |
| Ecuador | Other Lands | 3.10 | 3.13 |
|  | PAs | 1.30 | 1.55 |
|  | ITs | 1.39 | 2.25 |
|  | OAs | 1.48 | 1.80 |
| Peru | Other Lands | 6.57 | 6.72 |
|  | PAs | 10.15 | 11.70 |
|  | ITs | 1.94 | 3.24 |
|  | OAs | 6.37 | 5.62 |
| Brazil | Other Lands | 6.19 | 4.26 |
|  | PAs | 6.12 | 4.25 |
|  | ITs | 6.11 | 4.25 |
|  | OAs | 5.86 | 4.20 |

Considering the spatial distribution of matched observations, we performed geographic discontinuity designs to understand how carbon stocks varied spatially throughout the boundaries of ITs, OAs, and PAs in 2003 and 2016. The geographic discontinuity designs estimate spatial effects. That is, the mean differences of carbon stocks inside and outside these land tenures for various distances around their boundaries, after controlling for spatial location. Overall, the geographic discontinuity designs show that carbon stocks increase inside the boundaries in 2003 and 2016 (Fig 6). To explain away these effects, an unmeasured covariate would require a stronger effect than ITs, OAs, or PAs, especially as the distance increase from the boundaries, and above and beyond the covariates of spatial location (S3B Appendix). As discussed below, the geographic discontinuity designs reveal spatial and spatial-temporal patterns across ITs, OAs, and PAs.

**Fig 6.**
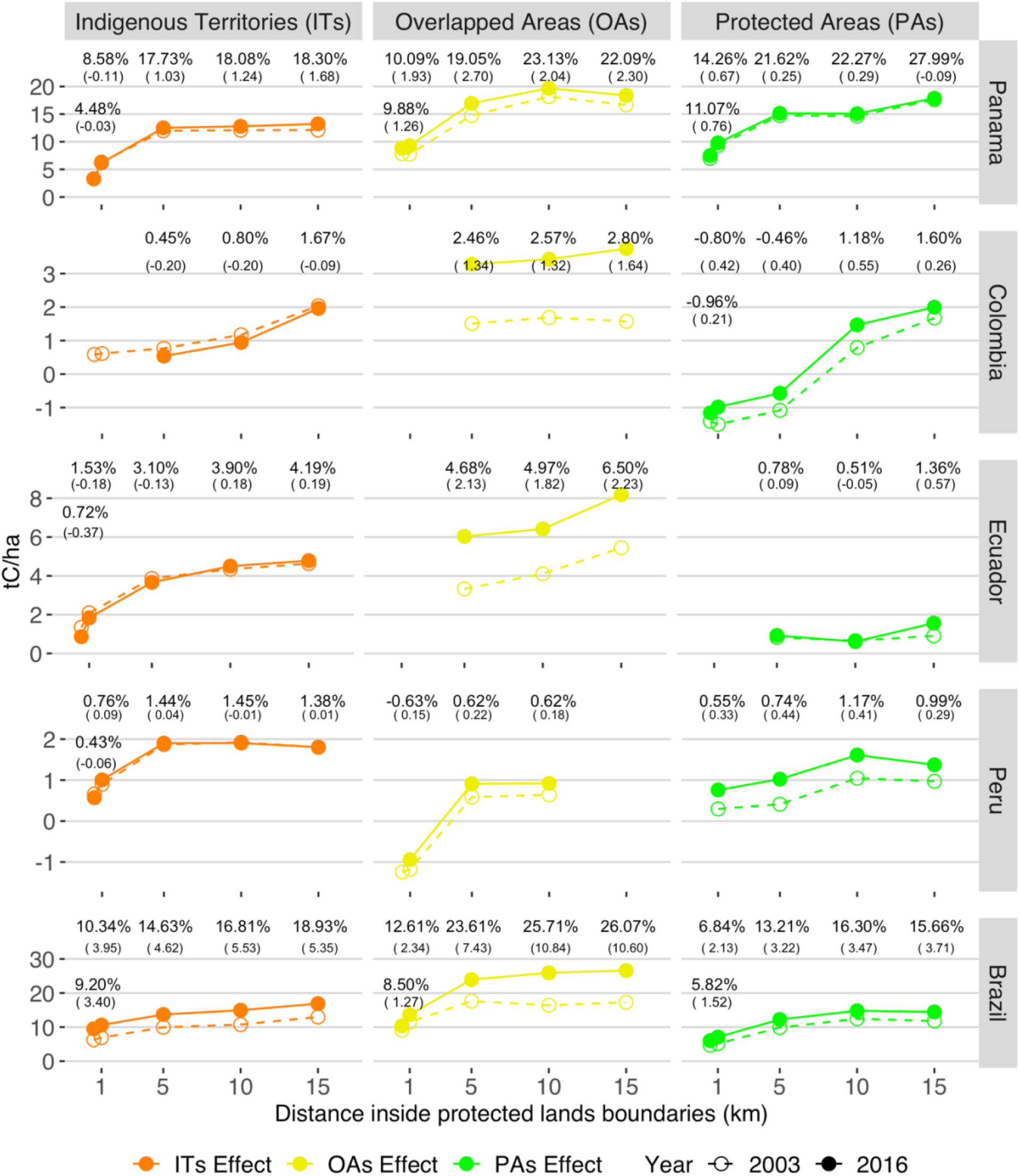
The spatial effect of ITs, OAs’, and PAs on carbon stocks during 2003 and 2016 in neotropical countries. Significant temporal effects (p < 0.05) are represented as points and percentages, indicating the additional/fewer carbon stocks secured inside the boundaries of ITs (orange), OAs (yellow), and PAs (green) relative to surrounding lands at multiple buffer distances (0-0.5 to 0-15 km). The spatial effects in 2003 are represented by empty points and dashed lines, while in 2016, they are full points and continuous lines. The values in parentheses represent the percentual increase/decrease in spatial effects between 2003 and 2016. Error bars reflect 95% confidence intervals for the temporal effects.

The spatial patterns of geographic discontinuity designs exhibit how ITs, OAs, and PAs influence carbon stocks within their boundaries. We found that the spatial effects of these land tenures tend to increase with the buffer distance to boundaries. At 0.5 km, the spatial effects are minimum or even insignificant; they become pronounced between 1 and 5 km and usually level off at 10km (Fig 6). For instance, ITs from Brazil in 2016 had carbon stocks 10.3% (21 t C/ha) larger than surrounding areas (102 t C/ha) when comparing a 1 km buffer. This spatial effect increased to 15% (27 t C/ha) at 5 km, 17% (∼30 t C/ha) at 10 km, and 19% (∼34 t C/ha) at 15 km. ITs in Panama and western Amazon Basin countries displayed a similar spatial effect. Except for Peru, OAs also had increasing spatial effects, and their influence on carbon stocks exceeded that of ITs and PAs. For example, OAs’ carbon stocks in Colombia did not differ from surrounding areas at 1km (120 t C/ha) in 2016 but had a spatial effect on carbon stocks of 2.5% (∼7 t C/ha) at 5 km, which is over five times higher than ITs’ and PAs’ effect in the same country. The spatial influence of PAs varied across countries. Relative to 10 km buffer comparisons, PAs spatial effects on carbon stocks reduce at 15 km in Brazil and Peru. At 1 and 5 km buffers, Colombia’s PAs had 0.80% and 0.46% fewer carbon stocks than surrounding lands, respectively. These resulting spatial patterns imply that allocating ITs and OAs generate boundaries that effectively conserve carbon stocks as PAs. Furthermore, the increasing effects on carbon stocks along with the distance to boundaries, more frequent in ITs and OAs, indicate that these land tenures shape forest landscapes by preserving the core and least accessible areas.

A spatial-temporal comparison of geographic discontinuities between 2003 and 2016 may indicate whether the boundaries of ITs, OAs, and PAs bring stability to carbon stocks. From 2003 to 2016, we found that the differences of carbon stocks inside and outside these land tenures increased, except for ITs in Colombia (Fig 6). Colombia’s ITs secured larger carbon stocks within their boundaries at 5 km and 10 km in 2016, but these spatial effects reduced 0.2%, potentially driven by a recovery in surrounding areas (S4D Appendix). The most substantial increases in spatial effects occurred among OAs. In Brazil, OAs’ spatial effect on carbon stocks increased by 11% (∼34 to 53 t C/ha) at 15 km in 2016, while ITs and PAs by 5.4% and 3.7%, respectively. Similarly, Ecuador’s OAs increased their spatial effects on carbon stocks by 2.2% at 15km, contrasting national PAs (0.6%) and ITs (0.2%). These increases between 2003 and 2016 in spatial effects suggest carbon stocks losses in surrounding areas that were buffered inside the boundaries of ITs, and PAs, but more prominently, in OAs.

## Discussion

In this study, we aim to estimate the temporal and spatial effects of allocating ITs, OAs and PAs, on carbon stocks across Neotropical Forests from Panama and the Amazon Basin. Considering that these land tenures tend to be located in higher elevations, steeper slopes, and greater distances to roads and human settlements than other lands, we control the effect of spatial location. Contrary to our hypothesis, ITs and OAs generally preserve carbon stocks and buffer losses as much as PAs. Over time, these land tenures secure more stable and higher carbon stocks than other lands between 2003 and 2016. Spatially, the geographic discontinuity designs show that carbon stocks increase inside the boundaries of ITs, OAs, and PAs. These temporal and spatial effects were conservative and had varied patterns across land tenures and countries.

### The effectiveness of Indigenous Territories in conserving forests and carbon stocks

Our findings highlight the need for a “spatially explicit” understanding of matching analysis regarding land tenure and forest conservation. Other studies have already incorporated “spatially explicit” methodologies. Gaveau et al. (22), for example, provides the spatial distribution of matched observation units among timber concessions, PAs and oil palm concessions in Kalimantan (Indonesia). Bowker et al. (20)in Africa and Zhao et al. (21) in China exclude from matching analysis other lands in a 10-km buffer around PAs. These studies attempt to avoid spatial autocorrelation by controlling sampling distance, while Negret et al. (58) test different post-matching models to control this bias and assess avoided deforestation in PAs from Colombia. Other studies use regression discontinuity designs to isolate some effects of spatial location and test the role of ITs and PAs boundaries (23,24). Our study presents an integrated approach. On the one hand, the temporal effect resembles matching methods that are not spatially explicit on sampled observation units (9,14,27,59). After exploring the spatial distribution of matched observation units, our findings point that they are located towards geographic boundaries, causing conservative estimates about ITs, OAs, and PAs. On the other hand, we use geographic discontinuity designs with matching analysis to directly control for spatial location and the geographic distance among observations, generating valid counterfactuals inside and outside these boundaries and maintaining conservative estimates (25). Hence, our study makes a novel methodological contribution to research by integrating matching analysis and geographic discontinuity designs to test the effectiveness of ITs’, OAs’, and PAs’ boundaries in conserving carbon stocks across neotropical countries.

Our findings support growing evidence indicating that decentralized forest governance can be effective in forest conservation (6,26,60,61). After controlling for spatial location and relative to other lands, we found that allocating indigenous lands (i.e., ITs and OAs) secured similar or even larger carbon stocks than PAs between 2003 and 2016 in Panama and Amazon Basin countries. These findings are in line with Nolte et al. (11), who showed that indigenous lands are more effective than PAs at curbing deforestation pressure in Brazil. By comparing indigenous lands (ITs and OAs) and PAs, our findings complement Blackman & Veit’s (14) estimates of avoided emissions from deforestation in ITs from Colombia and Brazil (14). However, they did not detect a discernible effect from Ecuador’s ITs, while our results estimated a positive effect on carbon stocks. Similarly, our results from Panama, where OAs had the most considerable effect on carbon stocks, partially contrast another study where PAs were the most effective in avoiding deforestation (12). These differences with previous studies might be attributable to our outcome variable (annual carbon stocks) that integrates deforestation, degradation, and recovery. Estimating carbon stock changes offer more accurate estimates regarding the effectiveness of ITs, OAs, and PAs, especially in countries where degradation emissions equal or exceed those from deforestation (e.g., Colombia, Ecuador, and Peru) (6). Thus, our results demonstrate that indigenous governance complements centralized forest governance, suggesting that titling ITs and formalizing shared governance in OAs while providing material and cultural benefits to their inhabitants can have a pivotal role in climate change mitigation.

Our geographic discontinuity designs provide conservative estimates regarding ITs’, OAs’, and PAs’ effect on carbon stocks within their boundaries. Although the assessments of PAs’ boundaries are common in the literature (31,62), they do not control for spatial location or compare different other land tenure categories. Our findings indicate that PAs’ carbon stocks are larger than surrounding areas, but these spatial effects vary within their boundaries. For example, PAs from Colombia seem only to avoid carbon stock losses 10 km inside their boundaries in 2003 and 2016. In Ecuador, PAs seem to have a stronger effect at 5 km than 10 km from their boundaries. These spatial patterns are not due to recent anthropogenic pressures and confirm the persistent inability of some PAs to reduce forest losses inside their boundaries (23,63,64). Furthermore, some anthropogenic pressures are not exclusively external, as our results show that a considerable amount of non-indigenous settlements is located in PAs.

Overall, the geographic discontinuities designs show that ITs and OAs tend to secure larger carbon stocks than their surroundings, and this difference tends to increase towards the least accessible areas. Similar results were found in ITs with granted property rights in Brazil (24) and titled IT’s in Colombia (23), which gradually decrease deforestation inside their boundaries. These gradual reductions in deforestation and degradation imply that indigenous land use decreases carbon stocks in the most accessible forests, where indigenous settlements tend to be located, and conserves core areas. Other studies have shown on a local scale these spatially limited impacts of indigenous land use, such as shifting agriculture and agroforestry, on carbon stocks (65,66). Additionally, our results reveal that the spatial effect of ITs and OAs remain temporarily stable similar to cases from Mexico and Ecuador (67,68). As established in the introduction, indigenous governance may explain this spatially and temporarily stable land use. Indigenous governance comprises institutions known to limit access based on cultural or social affiliations, and those with guaranteed access may develop and enforce rules that define the temporal and spatial extent of local livelihoods (69). Other factors, such as limited accessibility due to walking distances and changing river navigability, could also limit land use (70). Consequently, after controlling for spatial location, our results are among the first to establish that indigenous land use in neotropical forests may have a limited and stable spatial impact on carbon stocks.

### National contexts matter

Nonetheless, the current and future effects of allocating ITs, OAs, and PAs on carbon stocks are influenced by national contexts. General geographical trends indicate that these land tenures in Panama and Brazil have wider temporal and spatial effects on carbon stocks than Colombia, Ecuador, and Peru. These geographical differences reflect past trends of extensive forest loss in other lands from Panama (71) and Brazil (72). Moreover, the increasing differences in carbon stocks among ITs, OAs, and PAs with other lands after controlling for spatial location, highlight a growing pressure on neotropical forests. Consequently, their capacity to preserve or reduce carbon stock losses is likely to change. Between 2000 and 2013, tropical South America lost 7.3% of intact forest lands, mostly caused by the expansion of agriculture (73). PAs in Colombia are witnessing an increase in deforestation around their boundaries after the Peace Agreement with the Revolutionary Armed Forces (FARC) (64). ITs and PAs in southern Peru are threatened by growing road infrastructure, land invasions, illegal gold mining, and coca production (74). Oil blocks in the Ecuadorian Amazon will expand in cover from 32% to 68%, overlapping with biodiversity hotspots in PAs and ITs (75). In Brazil, limited law enforcement to prevent forest loss from soy, meat, and timber production in the Amazon Basin converge with recent setbacks in the land tenure security of ITs (76). Land invasions and deforestation in Panama also pose a threat to ITs (77). In this sense, as deforestation and degradation persist, countries’ climate benefits from forests are increasingly dependent on the stability of ITs’, OAs’, and PAs’ carbon stocks. The increasing dependence on stable forests points to the need to protect them through land use planning and resource allocation in institutions at the international, national, and sub-national levels (78,79).

### Study limitations

Finally, our study has some limitations. As with any estimates after matching analysis, the temporal and spatial effects are potentially biased by unmeasured covariates unrelated to the observed covariates (49). Nevertheless, the sensitivity analyses offer a transparent assessment regarding the influence required from unmeasured covariates to explain away our current estimates. The observed covariates still create a general classification of accessibility and forest loss pressures and control for spatial location. Despite using stratified sampling matching, known to effectively reduce covariate imbalances and the variability of treatment effects (e.g., temporal and spatial effects) (80), further research would benefit from comparing stratified and random sampling matching. We also aimed to identify the overall influence of ITs, OAs, and PAs across neotropical forests to solve the limited geographical scales and homogeneous pressures on forest loss in similar studies (32). However, these tenure categories represent different and diverse realities in each country. For instance, OAs in Colombia are subject to a policy that requires National Park Authorities to establish co-management agreements with Indigenous communities (81), which is not necessarily the case in other countries. Even in subnational scales, PAs comprise a broad spectrum of restrictions, local agents, and land use dynamics (26,82). Moreover, the geographic discontinuity designs and linear mixed models account for the influence of local or geographically restricted effects, but they do not incorporate the influence of external interventions such as REDD+ and payments for ecosystem services. Future studies could exploit the advantages of matching analysis and geographic discontinuity designs to explore the influence of PAs’ restrictions and external interventions. Finally, the outcome variable also brings limitations because it does not differentiate carbon stock losses due to deforestation and degradation; rather, it provides a comprehensive measure (i.e., aboveground carbon stocks) that captures forest conservation effectiveness beyond deforestation.

## Conclusions

After controlling the influence of spatial location, we found that ITs and OAs with decentralized forest governance represent effective natural climate solutions. Particularly, these indigenous lands and PAs have similar temporal and spatial effects on carbon stocks in Panama and Amazon Basin countries. Considering that the observation units sampled by matching are located along the boundaries of these land tenures, the temporal and spatial effects are conservative. Consequently, our findings show that indigenous peoples are supporting Nationally Determined Contributions (NDCs) under the Paris Agreement. Brazil and Ecuador expect to receive their first results-based payments from the Green Climate Fund corresponding to 96.5 and 18.6 million USD, respectively (83). For the critical role they play in reducing net carbon emissions, indigenous peoples must become recipients of such benefits, independent of the opportunity costs of avoided deforestation and degradation (84).

## Acknowledgements

For valuable comments and suggestions, we thank Frédéric Guichard, Margaret Kalacska, Divya Sharma, Paola Fajardo, Chris Madsen, and Matthieu Guillemette. Data analysis and storage was enabled in part by support provided by Calcul Québec (www.calculquebec.ca) and Compute Canada (www.computecanada.ca).

## Supporting information

**S1A Appendix. Geospatial Information and its sources by country.**

**S1B Appendix. Covariates mean differences between PAs, ITs, and OAs with other lands by country and their statistical significance from Mann Whitney U tests.**

*** p < 0.001, ** p < 0.01, * p < 0.05.

**S1C Appendix. Coarsening Choices applied through Coarsened Exact Matching (CEM) by country across PAs, ITs, and OAs.**

**S2A Appendix. Covariates standard mean differences between ITs, OAs, and PAs with other lands before (Pre-Match) and after matching analysis (Matched) across neotropical countries**.

**S2B Appendix. Kolmogorov-Smirnov statistics of covariates between ITs, OAs, and PAs with other lands before (Pre-Match) and after matching analysis (Matched) in neotropical countries**.

**S2C Appendix. Covariate standard mean differences before (Pre-Match) and after matching (Matched) in geographic discontinuity designs across the boundaries of ITs, OAs and PAs in neotropical countries**.

**S2D Appendix. Kolmogorov-Smirnov statistics of covariates before (Pre-Match) and after matching (Matched) in geographic discontinuity designs across the boundaries of ITs, OAs and PAs in neotropical countries**.

**S2E Appendix. Covariate falsification tests derived from linear mixed models in geographic discontinuity designs across the boundaries of ITs, OAs and PAs in neotropical countries**.

**S3A Appendix. Sensitivity analysis in the temporal effects of ITs, OAs, and PAs on carbon stocks in 2003 and 2016 across neotropical countries.** The temporal effect ratio is equivalent to the probability of a positive temporal effect in the treatment (i.e., ITs, OAs, and PAs) divided by the probability of a positive temporal effect in the control (other lands). The E-value represents the minimum strength that an unmeasured covariate would need to have with the treatment (i.e., ITs, OAs, PAs) and their temporal effect, for the treatments and temporal effect association not to be causal.

**S3B Appendix. Sensitivity analysis in the spatial effects of ITs’, OAs’, and PAs’ boundaries on carbon stocks in 2003 and 2016 across neotropical countries.** The spatial effect ratio is equivalent to the probability of a positive spatial effect in the treatment (i.e., ITs, OAs, and PAs) divided by the probability of a positive spatial effect in the control (other lands). The E-value represents the minimum strength that an unmeasured covariate would need to have with the treatment (i.e., ITs, OAs, PAs) and their spatial effect, for the treatments and spatial effect association not to be causal.

**S4A Appendix. The temporal effect of ITs, OAs, and PAs on carbon stocks between 2003 and 2016 across neotropical countries.** Each point are the significant annual effects (p < 0.05) of ITs (orange) OAs (yellow), and PAs (green) on carbon stocks. The temporal effects represent the annual differences of carbon stocks between ITs, OAs, and PAs with other lands after controlling for the spatial location through matching analysis and linear mixed models. Error bars reflect 95% confidence intervals for the temporal effect derived from the linear mixed models.

**S4B Appendix**. **The carbon stocks baseline of ITs’, OAs’, and PAs’ temporal effects across neotropical countries.** Each point represents the mean annual carbon stocks found in other lands (i.e., carbon stocks baseline) that share spatial location covariates with ITs (orange), OAs (yellow), and PAs (green) after matching analysis and linear mixed models. Error bars reflect 95% confidence intervals for the carbon stocks baselines derived from the linear mixed models.

**S4C Appendix**. **Distribution of human settlements inside the boundaries of ITs, OAs, and PAs across neotropical countries.** The human settlements included are those registered by national institutions in Amazon Basin countries and STRI in Panama. Data sources are shown in the S1 Appendix.

**S4D Appendix. The carbon stocks baseline outside the boundaries of ITs, OAs, and PAs across neotropical countries.** Full (2016) or empty (2003) points represent the mean annual carbon stocks found in other lands (i.e., carbon stocks baseline) outside the boundaries of ITs (orange), OAs (yellow), and PAs (green) at a certain buffer distance. Error bars reflect 95% confidence intervals for the carbon stocks baselines derived from linear mixed models.

## References

1. Griscom BW, Adams J, Ellis PW, Houghton RA, Lomax G, Miteva DA, et al. Natural climate solutions. Proc Natl Acad Sci [Internet]. 2017;114(44):11645–50. Available from: http://www.pnas.org/lookup/doi/10.1073/pnas.1710465114

2. MacKinnon K, Dudley N, Sandwith T. Natural solutions: protected areas helping people to cope with climate change. Oryx [Internet]. 2011 Oct 8;45(4):461–2. Available from: https://www.cambridge.org/core/product/identifier/S0030605311001608/type/journal_article

3. Morales-Hidalgo D, Oswalt SN, Somanathan E. Status and trends in global primary forest, protected areas, and areas designated for conservation of biodiversity from the Global Forest Resources Assessment 2015. For Ecol Manage. 2015;352:68–77.

4. Keenan RJ, Reams GA, Achard F, de Freitas J V., Grainger A, Lindquist E. Dynamics of global forest area: Results from the FAO Global Forest Resources Assessment 2015. For Ecol Manage. 2015;352:9–20.

5. Garnett ST, Burgess ND, Fa JE, Fernández-Llamazares Á, Molnár Z, Robinson CJ, et al. A spatial overview of the global importance of Indigenous lands for conservation. Nat Sustain [Internet]. 2018;1(7):369–74. Available from: http://dx.doi.org/10.1038/s41893-018-0100-6

6. Walker WS, Gorelik SR, Baccini A, Aragon-Osejo JL, Josse C, Meyer C, et al. The role of forest conversion, degradation, and disturbance in the carbon dynamics of Amazon indigenous territories and protected areas. Proc Natl Acad Sci [Internet]. 2020 Feb 11;117(6):3015–25. Available from: http://www.pnas.org/lookup/doi/10.1073/pnas.1913321117

7. Joppa LN, Pfaff A. High and far: Biases in the location of protected areas. PLoS One. 2009;4(12):1–6.

8. Joppa LN, Pfaff A. Reassessing the forest impacts of protection: The challenge of nonrandom location and a corrective method. Ann N Y Acad Sci. 2010;1185:135–49.

9. Nelson A, Chomitz KM. Effectiveness of strict vs. multiple use protected areas in reducing tropical forest fires: A global analysis using matching methods. PLoS One. 2011;6(8).

10. Andam KS, Ferraro PJ, Pfaff A, Sanchez-Azofeifa GA, Robalino JA. Measuring the effectiveness of protected area networks in reducing deforestation. Proc Natl Acad Sci U S A. 2008;105(42):16089–94.

11. Nolte C, Agrawal A, Silvius KM, Britaldo SSF. Governance regime and location influence avoided deforestation success of protected areas in the Brazilian Amazon. Proc Natl Acad Sci U S A. 2013;110(13):4956–61.

12. Vergara-Asenjo G, Potvin C. Forest protection and tenure status: THE key role of indigenous peoples and protected areas in Panama. Glob Environ Chang. 2014;28(1):205–15.

13. Anderson CM, Asner GP, Llactayo W, Lambin EF. Overlapping land allocations reduce deforestation in Peru. Land use policy. 2018;79(August 2017):174–8.

14. Blackman A, Veit P. Titled Amazon Indigenous Communities Cut Forest Carbon Emissions. Ecol Econ [Internet]. 2018;153(July 2017):56–67. Available from: https://doi.org/10.1016/j.ecolecon.2018.06.016

15. Mertz O, Müller D, Sikor T, Hett C, Heinimann A, Castella JC, et al. The forgotten D: Challenges of addressing forest degradation in complex mosaic landscapes under REDD+. Geogr Tidsskr. 2012;112(1):63–76.

16. van Vliet N, Mertz O, Heinimann A, Langanke T, Pascual U, Schmook B, et al. Trends, drivers and impacts of changes in swidden cultivation in tropical forest-agriculture frontiers: A global assessment. Glob Environ Chang [Internet]. 2012;22(2):418–29. Available from: http://dx.doi.org/10.1016/j.gloenvcha.2011.10.009

17. Bruun TB, de Neergaard A, Lawrence D, Ziegler AD. Environmental consequences of the demise in Swidden cultivation in Southeast Asia: Carbon storage and soil quality. Hum Ecol. 2009;37(3):375–88.

18. Andrade RB, Balch JK, Parsons AL, Armenteras D, Roman-Cuesta RM, Bulkan J. Scenarios in tropical forest degradation: Carbon stock trajectories for REDD+. Carbon Balance Manag. 2017;12(1):1–7.

19. Karsenty A, Romero C, Cerutti PO, Doucet JL, Putz FE, Bernard C, et al. Deforestation and timber production in Congo after implementation of sustainable management policy: A reaction to the article by J.S. Brandt, C. Nolte and A. Agrawal (Land Use Policy 52:15–22). Land use policy [Internet]. 2017;65:62–5. Available from: http://dx.doi.org/10.1016/j.landusepol.2017.02.032

20. Bowker JN, De Vos A, Ament JM, Cumming GS. Effectiveness of Africa’s tropical protected areas for maintaining forest cover. Conserv Biol. 2017;31(3):559–69.

21. Zhao H, Wu R, Long Y, Hu J, Yang F, Jin T, et al. Individual-level performance of nature reserves in forest protection and the effects of management level and establishment age. Biol Conserv [Internet]. 2019;233(February):23–30. Available from: https://doi.org/10.1016/j.biocon.2019.02.024

22. Gaveau DLA, Kshatriya M, Sheil D, Sloan S, Molidena E, Wijaya A, et al. Reconciling Forest Conservation and Logging in Indonesian Borneo. PLoS One. 2013;8(8).

23. Bonilla-Mejía L, Higuera-Mendieta I. Protected Areas under Weak Institutions: Evidence from Colombia. World Dev [Internet]. 2019 Oct;122:585–96. Available from: https://linkinghub.elsevier.com/retrieve/pii/S0305750X19301718

24. Baragwanath K, Bayi E. Collective property rights reduce deforestation in the Brazilian Amazon. Proc Natl Acad Sci U S A. 2020;117(34):20495–502.

25. Keele LJ, Titiunik R, Zubizarreta JR. Enhancing a geographic regression discontinuity design through matching to estimate the effect of ballot initiatives on voter turnout. J R Stat Soc Ser A Stat Soc. 2015;178(1):223–39.

26. Blackman A. Strict versus mixed-use protected areas: Guatemala’s Maya Biosphere Reserve. Ecol Econ [Internet]. 2015;112:14–24. Available from: http://dx.doi.org/10.1016/j.ecolecon.2015.01.009

27. Pfaff A, Robalino J, Lima E, Sandoval C, Herrera LD. Governance, Location and Avoided Deforestation from Protected Areas: Greater Restrictions Can Have Lower Impact, Due to Differences in Location. World Dev. 2014;55:7–20.

28. Ferretti-Gallon K, Busch J. What Drives Deforestation and What Stops it? A Meta-Analysis of Spatially Explicit Econometric Studies. SSRN Electron J [Internet]. 2014;(April 2014). Available from: http://www.ssrn.com/abstract=2458040

29. Geist HJ, Lambin EF. Proximate Causes and Underlying Driving Forces of Tropical Deforestation. Bioscience. 2002;52(2):143.

30. Barber CP, Cochrane MA, Souza CM, Laurance WF. Roads, deforestation, and the mitigating effect of protected areas in the Amazon. Biol Conserv [Internet]. 2014 Sep;177(2014):203–9. Available from: http://dx.doi.org/10.1016/j.biocon.2014.07.004

31. Joppa LN, Loarie SR, Pimm SL. On the protection of “protected areas.” Proc Natl Acad Sci U S A. 2008;105(18):6673–8.

32. Börner J, Schulz D, Wunder S, Pfaff A. The effectiveness of forest conservation policies and programs. Annu Rev Resour Econ. 2020;12:45–64.

33. Villalba U. Buen Vivir vs Development: A paradigm shift in the Andes? Third World Q. 2013;34(8):1427–42.

34. Walsh C. Development as Buen Vivir: Institutional arrangements and (de)colonial entanglements. Development. 2010;53(1):15–21.

35. Berkes F, Colding J, Folke C. Rediscovery of Traditional Ecological Knowledge as Adaptive Management. Ecol Appl [Internet]. 2000;10(5):1251. Available from: http://search.ebscohost.com/login.aspx?direct=true&db=edsjaa&AN=jstor.10.2307.2641280&lang=es&site=eds-live

36. Hosonuma N, Herold M, De Sy V, De Fries RS, Brockhaus M, Verchot L, et al. An assessment of deforestation and forest degradation drivers in developing countries. Environ Res Lett. 2012;7(4).

37. Börner J, Vosti SA. Managing Tropical Forest Ecosystem Services: An Overview of Options. In 2013. p. 21–46. Available from: http://link.springer.com/10.1007/978-94-007-5176-7_2

38. Chhatre A, Agrawal A. Trade-offs and synergies between carbon storage and livelihood benefits from forest commons. Proc Natl Acad Sci [Internet]. 2009;106(42):17667–70. Available from: http://www.pnas.org/cgi/doi/10.1073/pnas.0905308106

39. de Koning F, Aguiñaga M, Bravo M, Chiu M, Lascano M, Lozada T, et al. Bridging the gap between forest conservation and poverty alleviation: the Ecuadorian Socio Bosque program. Environ Sci Policy [Internet]. 2011 Aug;14(5):531–42. Available from: https://linkinghub.elsevier.com/retrieve/pii/S1462901111000657

40. Sills EO, Atmadja SS, de Sassi C, Duchelle AE, Kweka D, Resosudarmo IAP, et al. REDD+ on the ground: A case book of subnational initiatives across the globe [Internet]. REDD+ on the ground: A case book of subnational initiatives across the globe. Bogor: Center for International Forestry Research (CIFOR); 2014. 2014 p. Available from: http://www.cifor.org/library/5202/redd-on-the-ground-a-case-book-of-subnational-initiatives-across-the-globe

41. UNEP-WCMC, IUCN, NGS. Protected Planet Report 2018. Tracking progress towards global targets for protected areas [Internet]. UNEP-WCMC, IUCN, NGS, editors. Cambridge UK; Gland, Switzerland; and Washington, D.C., USA.; 2018 [cited 2019 Jun 12]. Available from: www.unep-wcmc.org

42. Borrini-Feyerabend G, Pimbert M, Farvar T, Kothari A, Renard Y, Farvar MT, et al. Sharing power: Learning-by-doing in co-management of natural resources throughout the world. [Internet]. Cenesta, Teheran: IIED and IUCN; 2004. 456 p. Available from: http://www.iucn.org/about/union/commissions/ceesp/ceesp_publications/sharing_power.cfm#sp_contents

43. Baccini A, Goetz SJ, Walker WS, Laporte NT, Sun M, Sulla-Menashe D, et al. Estimated carbon dioxide emissions from tropical deforestation improved by carbon-density maps. Nat Clim Chang [Internet]. 2012;2(3):182–5. Available from: http://dx.doi.org/10.1038/nclimate1354

44. Baccini A, Walker W, Carvalho L, Farina M, Sulla-Menashe D, Houghton RA. Tropical forests are a net carbon source based on aboveground measurements of gain and loss. Supplementary Materials. Science (80-) [Internet]. 2017 Oct 13;358(6360):230–4. Available from: http://www.sciencemag.org/lookup/doi/10.1126/science.aam5962

45. Diamond A, Sekhon J. Genetic Matching for Estimating Causal Effects. Rev Econ Stat [Internet]. 2012;95(July):932–45. Available from: http://sekhon.polisci.berkeley.edu/papers/GenMatch.pdf

46. Iacus SM, King G, Porro G. cem : Software for Coarsened Exact Matching. J Stat Softw. 2015;30(9).

47. Ho DE, Imai K, King G, Stuart EA. MatchIt : Nonparametric Preprocessing for Parametric Causal Inference. J Stat Softw. 2015;42(8).

48. Iacus SM, King G, Porro G. Society for Political Methodology Causal Inference without Balance Checking: Coarsened Exact Matching [Internet]. Vol. 20, Source: Political Analysis. 2012 [cited 2019 May 1]. Available from: http://gking.harvard.edu/cem;

49. Stuart EA. Matching methods for causal inference: A review and a look forward. Stat Sci. 2010;25(1):1–21.

50. Greifer N. Package ‘cobalt’ [Internet]. 2021. Available from: https://cran.r-project.org/web/packages/cobalt/cobalt.pdf

51. Bates D, Maechler M, Bolker B, Walker S, Chistensen RHB, Singman H, et al. Linear mixed-effects models using “Eigen” and S4. 2019.

52. Keele LJ, Titiunik R. Geographic boundaries as regression discontinuities. Polit Anal. 2015;23(1):127–55.

53. Angelsen A. Policies for reduced deforestation and their impact on agricultural production. Proc Natl Acad Sci U S A. 2010;107(46):19639–44.

54. Liu W, Kuramoto SJ, Stuart EA. An Introduction to Sensitivity Analysis for Unobserved Confounding in Nonexperimental Prevention Research. Prev Sci [Internet]. 2013 Dec 14;14(6):570–80. Available from: https://www.ncbi.nlm.nih.gov/pmc/articles/PMC3624763/pdf/nihms412728.pdf

55. VanderWeele TJ, Ding P. Sensitivity Analysis in Observational Research: Introducing the E-Value. Ann Intern Med [Internet]. 2017 Aug 15;167(4):268. Available from: http://annals.org/article.aspx?doi=10.7326/M16-2607

56. VanderWeele TJ, Ding P, Mathur M. Technical Considerations in the Use of the E-Value. J Causal Inference [Internet]. 2019 Sep 25;7(2):1–11. Available from: https://www.degruyter.com/document/doi/10.1515/jci-2018-0007/html

57. Mathur MB, Smith LH, Peng D, VanderWeele TJ. Package ‘EValue’ [Internet]. 2021. Available from: https://cran.r-project.org/web/packages/EValue/EValue.pdf

58. Negret PJ, Di-Marco M, Sonter LJ, Rhodes J, Possingham HP, Maron M. Effects of spatial autocorrelation and sampling design on estimates of protected area effectiveness. Conserv Biol [Internet]. 2020 Apr 28;cobi.13522. Available from: https://onlinelibrary.wiley.com/doi/abs/10.1111/cobi.13522

59. Vergara-asenjo G, Sharma D, Potvin C. Engaging Stakeholders : Assessing Accuracy of Participatory Mapping of Land Cover in Panama. 2015;8(December):432–9.

60. Porter-Bolland L, Ellis EA, Guariguata MR, Ruiz-Mallén I, Negrete-Yankelevich S, Reyes-García V. Community managed forests and forest protected areas: An assessment of their conservation effectiveness across the tropics. For Ecol Manage [Internet]. 2012;268:6–17. Available from: http://dx.doi.org/10.1016/j.foreco.2011.05.034

61. Newton P, A Oldekop J, Brodnig G, Karna BK, Agrawal A. Carbon, biodiversity, and livelihoods in forest commons: synergies, trade-offs, and implications for REDD+. Environ Res Lett [Internet]. 2016;11(4):1–7. Available from: http://dx.doi.org/10.1088/1748-9326/11/4/044017

62. Spracklen BD, Kalamandeen M, Galbraith D, Gloor E, Spracklen D V. A global analysis of deforestation in moist tropical forest protected areas. PLoS One. 2015;10(12):1–16.

63. Lui G V., Coomes DA. Tropical nature reserves are losing their buffer zones, but leakage is not to blame. Environ Res [Internet]. 2016;147:580–9. Available from: http://dx.doi.org/10.1016/j.envres.2015.11.008

64. Clerici N, Armenteras D, Kareiva P, Botero R, Ramírez-Delgado JP, Forero-Medina G, et al. Deforestation in Colombian protected areas increased during post-conflict periods. Sci Rep. 2020;10(1):1–10.

65. Kirby KR, Potvin C. Variation in carbon storage among tree species: Implications for the management of a small-scale carbon sink project. For Ecol Manage. 2007;246(2–3):208–21.

66. Tschakert P, Coomes OT, Potvin C. Indigenous livelihoods, slash-and-burn agriculture, and carbon stocks in Eastern Panama. Ecol Econ. 2007;60(4):807–20.

67. Puc-Alcocer M, Arce-Ibarra AM, Cortina-Villar S, Estrada-Lugo EIJ. Rainforest conservation in Mexico’s lowland Maya area: Integrating local meanings of conservation and land-use dynamics. For Ecol Manage [Internet]. 2019;448(June):300–11. Available from: https://doi.org/10.1016/j.foreco.2019.06.016

68. Gray C, Bilsborrow R. Stability and change within indigenous land use in the Ecuadorian Amazon. Glob Environ Chang [Internet]. 2020 Jul;63(August 2019):102116. Available from: https://doi.org/10.1016/j.gloenvcha.2020.102116

69. Berkes F. Sacred Ecology. Second Edi. New York: Routledge, Taylor and Francis Group; 2008.

70. Jakovac CC, Dutrieux LP, Siti L, Peña-Claros M, Bongers F. Spatial and temporal dynamics of shifting cultivation in the middle-Amazonas river: Expansion and intensification. PLoS One. 2017;12(7):1–15.

71. Redo DJ, Grau HR, Aide TM, Clark ML. Asymmetric forest transition driven by the interaction of socioeconomic development and environmental heterogeneity in Central America. Proc Natl Acad Sci [Internet]. 2012;109(23):8839–44. Available from: http://www.pnas.org/cgi/doi/10.1073/pnas.1201664109

72. Fearnside P, Fearnside P. Deforestation of the Brazilian Amazon. Oxford Research Encyclopedia of Environmental Science. 2017. 1–52 p.

73. Potapov P, Hansen MC, Laestadius L, Turubanova S, Yaroshenko A, Thies C, et al. The last frontiers of wilderness: Tracking loss of intact forest landscapes from 2000 to 2013. Sci Adv. 2017;3(1):1–14.

74. Gallice GR, Larrea-Gallegos G, Vázquez-Rowe I. The threat of road expansion in the Peruvian Amazon. Oryx [Internet]. 2019 Apr 27;53(2):284–92. Available from: https://www.cambridge.org/core/product/identifier/S0030605317000412/type/journal_article

75. Lessmann J, Fajardo J, Muñoz J, Bonaccorso E. Large expansion of oil industry in the Ecuadorian Amazon: biodiversity vulnerability and conservation alternatives. Ecol Evol. 2016;6(14):4997–5012.

76. Carvalho WD, Mustin K, Hilário RR, Vasconcelos IM, Eilers V, Fearnside PM. Deforestation control in the Brazilian Amazon: A conservation struggle being lost as agreements and regulations are subverted and bypassed. Perspect Ecol Conserv [Internet]. 2019 Jul;17(3):122–30. Available from: https://linkinghub.elsevier.com/retrieve/pii/S2530064418301263

77. Vergara-Asenjo G, Mateo-Vega J, Alvarado A, Potvin C. A participatory approach to elucidate the consequences of land invasions on REDD+ initiatives: A case study with Indigenous communities in Panama. PLoS One [Internet]. 2017;12(12):1–19. Available from: http://dx.doi.org/10.1371/journal.pone.0189463

78. Busch J, Amarjargal O. Authority of Second-Tier Governments to Reduce Deforestation in 30 Tropical Countries. Front For Glob Chang. 2020;3(January):1–14.

79. Funk JM, Aguilar-Amuchastegui N, Baldwin-Cantello W, Busch J, Chuvasov E, Evans T, et al. Securing the climate benefits of stable forests. Clim Policy. 2019;19(7):845–60.

80. Iacus SM, King G, Porro G. A Theory of Statistical Inference for Matching Methods in Causal Research. Polit Anal [Internet]. 2019 Jan 4;27(1):46–68. Available from: https://www.cambridge.org/core/product/identifier/S1047198718000293/type/journal_article

81. Unidad Administrativa del Sistema de Parques Nacionales Naturales. Política de Participación Social en la Conservación [Internet]. Bogotá: UAESPNN; 2001. 83 p. Available from: http://www.parquesnacionales.gov.co/PNN/portel/libreria/pdf/politicadeparticipacinsocial2.pdf

82. Radachowsky J, Ramos VH, McNab R, Baur EH, Kazakov N. Forest concessions in the Maya Biosphere Reserve, Guatemala: A decade later. For Ecol Manage. 2012;268:18–28.

83. Maniatis D, Scriven J, Jonckheere I, Laughlin J, Todd K. Toward REDD+ Implementation. Annu Rev Environ Resour. 2019;44(1):373–98.

84. Karsenty A, Vogel A, Castell F. “Carbon rights”, REDD+ and payments for environmental services. Environ Sci Policy [Internet]. 2014 Jan;35:20–9. Available from: https://linkinghub.elsevier.com/retrieve/pii/S1462901112001463

